# Differential regulation of developmental stages supports a linear model for *C. elegans* postembryonic development

**DOI:** 10.1101/2021.05.03.442510

**Authors:** Alejandro Mata-Cabana, Francisco Javier Romero-Expósito, Mirjam Geibel, Francine Amaral Piubeli, Martha Merrow, María Olmedo

## Abstract

The repetitive nature of *C. elegans* postembryonic development is considered an oscillatory process, a concept that has gained traction from regulation by a circadian clock gene homolog. Nevertheless, each larval stage has a defined duration and entails specific events. We have measured the duration of each stage of development for over 2,500 larvae, under varied environmental conditions known to alter overall developmental rate. Our results show that distinct developmental stages respond differently to environmental perturbations, including changes in temperature, in food quantity and quality, and amount of insulin signaling. Furthermore, our high-resolution measurement of the effect of temperature on the stage-specific duration of development has unveiled novel features of temperature dependence in *C. elegans* postembryonic development. Altogether, our results support a model of linear progression of *C. elegans* development, with the duration of each stage determined by a unique program.

## INTRODUCTION

*C. elegans* postembryonic development consists of four larval stages (L1 to L4). At the beginning of each larval stage, a nutritional checkpoint controls initiation (Schindler et al., 2014). Towards the end of each larval stage, molting occurs. The initiation of each molt is marked by the sealing of the buccal cavity of the larva, which impedes food intake (Singh and Sulston, 1978). During the molts, larvae enter a quiescent sleep-like state (Raizen et al., 2008). The end of each molt is determined by the removal of the old cuticle, called ecdysis, thus concluding that the larval stage. Molts and intermolts are therefore fundamentally different processes within each larval stage. Unfortunately, most measurements of postembryonic development do not allow precise resolution of the timing of molts and intermolts.

The repetitive process of molting is coupled to oscillatory expression of ~3,700 genes (Hendriks et al., 2014; Kim et al., 2013; Meeuse et al., 2020). One of the oscillating genes is *lin-42*, the homologue of the *Drosophila melanogaster* clock protein *period* (Jeon et al., 1999). LIN-42 has a reiterative function during larval development that is reminiscent of the cyclic functions of clock proteins (Monsalve et al., 2011). On this basis, *C. elegans* larval development has been compared to other rhythmic processes, especially daily or circadian rhythms (Meeuse et al., 2020; Monsalve et al., 2011; Monsalve and Frand, 2012; reviewed in Olmedo et al., 2017). However, the four larval stages have non-identical durations (Byerly et al., 1976; Faerberg et al., 2021; Meeuse et al., 2020; Monsalve et al., 2011; Olmedo et al., 2015) and are characterized by stage-specific patterns of cell division (Sulston and Horvitz, 1977). The stage specific fates are controlled by the heterochronic gene pathway. Heterochronic mutants show alterations in the normal order of stage-specific cell division patterns throughout the animal (Ambros and Horvitz, 1984). These mutants showed that the stage-specific patterns of cell division are modular (Rougvie and Moss, 2013). At the level of transcription, each stage is also modular. Once transcription is activated at each larval stage, it progresses until the next checkpoint. The transcriptional oscillation is therefore reset at the beginning of each larval stage (Meeuse et al., 2020; Stec et al., 2021).

A central question remains as to how the overall speed of postembryonic development is modulated by rate-limiting interventions. Although the progression of development is under strict genetic control (Rougvie and Moss, 2013), the speed of the process depends on a variety of environmental factors. If developmental rate is controlled by an oscillator, rate-limiting interventions that impact this oscillator would have similar effects in all stages of development, yielding slower or faster animals, with larval stages that scale proportionally to the duration of total development. This would resemble how zeitgebers are known to act on a circadian clock, even in non-natural constant conditions (Abraham et al., 2006; Aschoff, 1960). In contrast, if development proceeds in a linear manner, based on completing stage-specific events, perturbations that affect overall duration of development may have different impact on different stages, depending on the specific events that take place during each stage. The solution to this question calls for precise quantification of postembryonic development in response to varied environmental perturbations.

The speed of both embryonic and postembryonic development in *C. elegans* is temperature sensitive, as in many poikilothermic organisms. The published relationships show that postembryonic development can be completed in about 39 h at 25 °C compared with 75 h at 16 °C (Byerly et al., 1976). Changes in the bacterial diet and in insulin signaling also affect developmental rate (MacNeil et al., 2013; Ruaud et al., 2011; Shtonda and Avery, 2006; Uppaluri and Brangwynne, 2015). Both of these interventions may increase or decrease metabolic rates and available energy thus liberating more or less energy that can be applied to development. Although temperature, diet and insulin all impact metabolism, the presumably do so via distinct pathways: altered biochemical rates of reactions due to temperature, altered nutrient availability or energy metabolism as regulated by cell signaling.

In this work we have imposed environmental and genetic perturbations to characterize the stage-specific response of *C. elegans* postembryonic development using a quantitative high-throughput method. We quantified development of ~2,500 individually assayed larvae. Importantly, we monitored development continuously, with a time resolution of five minutes, and resolved the transitions between molts and intermolts. We analyzed interindividual variability of development, its dependence on temperature and food, and its regulation by insulin signaling. In all cases, we observe that interventions that alter developmental timing have a differential effect on the discrete stages of larval development. Furthermore, modification of the cell division pattern by downregulation of heterochronic genes suggests that the duration of the stages is linked to the divisions taking place during each stage. These observations support a mechanism based on a linear progression of development, where the events that take place in each stage are uniquely impacted by environmental perturbations.

## RESULTS AND DISCUSSION

### Quantitative analysis of *C. elegans* postembryonic development

We quantified *C. elegans* postembryonic developmental progression to evaluate interindividual variability. Using a luminometry-based method, we determined the length of larval stages. Unlike other methods, here the duration of the molting period is measured (Olmedo et al., 2015). As published elsewhere (Meeuse et al., 2020), we use a nomenclature that divides each larval stage into molt and intermolt (Fig. 1A). We thus used the measurements to determine the duration of each of the four complete larval stages (L1, L2, L3 and L4), as well as that of the intermolt and molt (*i.e*. L1=I1+M1).

**Figure 1.**
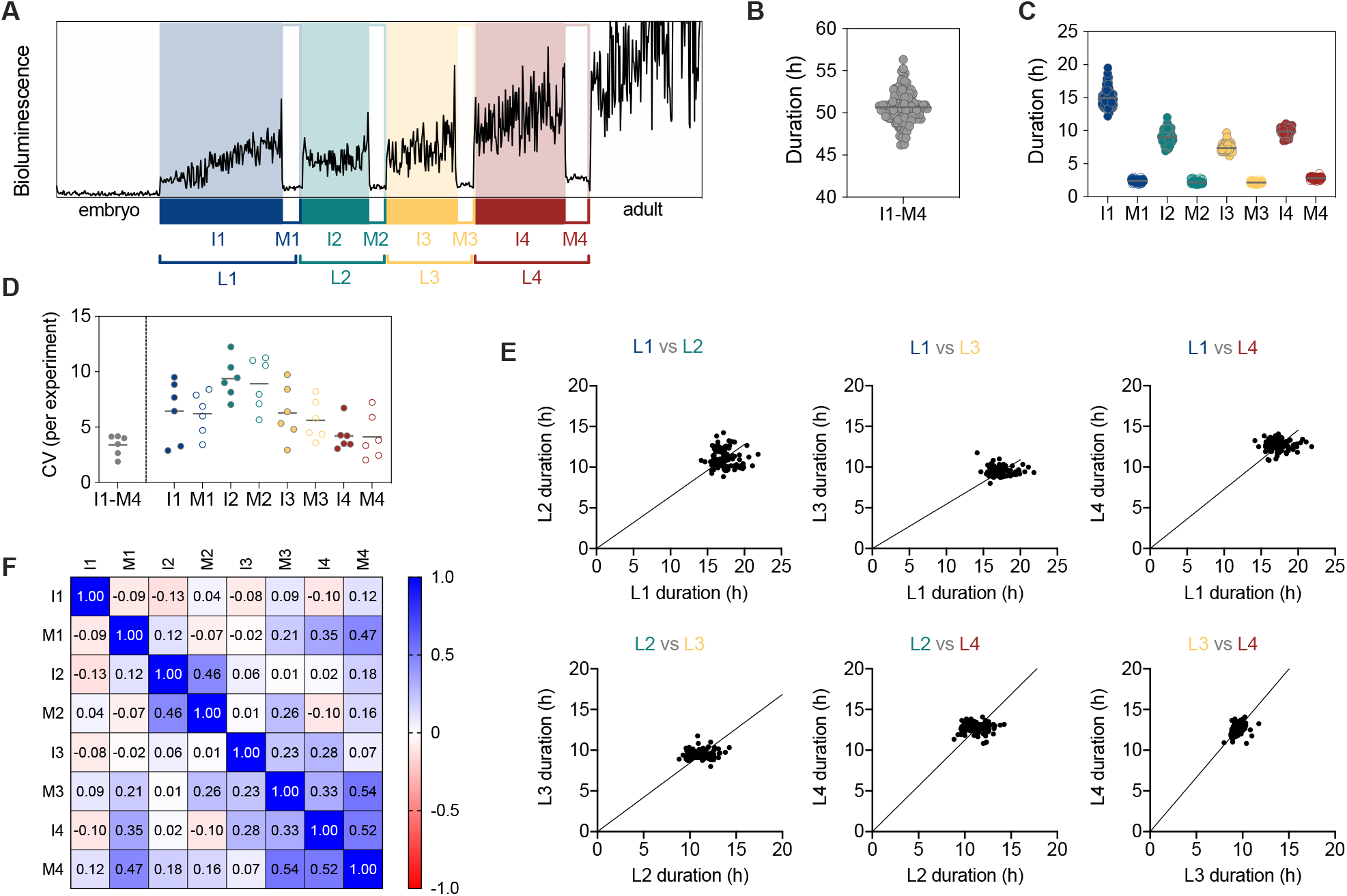
Quantitative analysis of development in 103 individual larvae. (A) Nomenclature of the different stages of development as defined by the luminometry assay. (B) Total duration of development, from hatching to adult ecdysis (I1-M4) for 103 larvae at 20 °C. (C) Duration of each stage of development for the larvae in (B). (D) Coefficient of variation for the complete development (I1-M4) and for each stage independently. Each dot represents the CV of an independent experiment. (E) Comparison of duration for the six possible pairs of larval stages. The line represents the linear regression of the data forced through the origin. (F) Pairwise correlation matrix for all combinations of developmental stages.

The average duration of development from hatching to adulthood for 103 larvae measured in six independent experiments at 20 °C was 50.67±1.95 hours (Fig. 1B). This is similar to previous observations at the same temperature (Byerly et al., 1976; Hirsh et al., 1976; Keil et al., 2016). The duration of each stage of postembryonic development is shown in figure 1C and Supplementary figure 1A. The coefficient of variation (CV) in the duration of the complete postembryonic development (from hatching to the end of M4) was 3.8%, lower than the 4.6% value recently measured by direct observation of movement of the N2 strain (Faerberg et al., 2021).

We also calculated the fraction of development in each larval stage by dividing the duration of each stage by the total duration of development. For the four larval stages (L1 to L4) we obtained the average values of 0.34, 0.22, 0.19 and 0.25 respectively (Supplementary Figs. 1B-C, left panels). These values are similar to those observed in multiple studies at different temperatures (Byerly et al., 1976; Faerberg et al., 2021; Gritti et al., 2016; Hirsh et al., 1976; Raizen et al., 2008). This means that the transgenic animals used for the luminometry assay conform to reported temporal structures of developmental timing of the wild-type strain, considered for both duration and variability.

Stage-by-stage analysis revealed that temporal precision differs between stages (Fig. 1D). The coefficient of variation is larger at the beginning of larval development, specially at L2, and lower at later stages, specially at L4. Furthermore, the coefficient of variation of the fractional duration showed a modest reduction compared to that of the total duration of each stage (Supplementary Fig. 1 B-C, right panels), as has been observed elsewhere (Faerberg et al., 2021). The observation that total larval development has a lower variability than individual stages suggests that the durations of larval stages do not scale proportionally across individuals. If some animals had consistently slower or faster development, interindividual variability should be similar between stages and similar to that of the complete process, which is not the case in our experiments (Fig. 1D). Instead, if there were negative correlations between the duration of stages, that is, if longer stages were compensated by a shorter duration in following stages, the overall variability would be reduced. To evaluate whether different developmental periods scale proportionally, we plotted the duration of the larval stages against each other for all six possible combinations. Most combinations yield results that suggested a lack of proportionality between the duration of larval stages (Fig. 1E). When we tested the correlation between each stage of larval development, we observed a mild negative correlation between I1 and each of the following intermolts. Although the coefficients of correlation are generally low, we observed a trend from negative correlations at the beginning of development to positive correlations when comparing late larval stages, especially between the third and fourth intermolts and molts (Fig. 1F). This contrasts with other studies which found proportionality (positive correlation) between the different stages of development (Faerberg et al., 2021; Filina et al., 2020). We tried to understand why we found a positive correlation only between the duration of late larval stages. These stages also show less variability than L1 and L2. Increased variability of L1 and L2 may be breaking the proportionality between stages. Since the larger variability of L1 and L2 has been recently observed using the N2 strain (Faerberg et al., 2021), we are inclined to think that this is a feature of *C. elegans* postembryonic development. Interindividual variability of developmental rate has been linked to maternal age (Perez et al., 2017). Indeed, when we analyzed stage by stage development of larva from mothers of different age, we observed that maternal age mainly impacts the duration of I1 and I2 (Supplementary Fig. 1D). Furthermore, maternal exposure to pheromones specifically delays I1 and molts of the progeny (Perez et al., 2020). Similar to these two examples, other factors that generate interindividual variability might impact development in a stage specific manner. Interindividual variability in the extension of specific stages would thus cause departures from proportionality.

We would also like to reflect on the possibility that proportionality would emerge from the combination of measurement of development from larvae raised at slightly different temperatures. Small variations in incubation temperature, as low as 0.85 °C, increases variability in postembryonic development comparable to that found in other studies (Supplementary Fig. 1E). Since the effect of temperature is similar in the different stages (especially when lacking resolution between molts and intermolts; see below), these small variations in temperature increase the correlation between the duration of stages. As an example, proportionality increased when we combined the results from the original 103 larvae measured at 20 °C (T20) with a group of 17 larvae (T20.85) that moved the average duration of development by ~30 min. The average duration of the T20.85 group could be obtained by an increase in experimental temperature of only 0.85 °C (as calculated by the temperature dependence of developmental rate, see below). When we added 18 more larvae (T21.25), the average duration of development changed by ~1 hour. The average duration of the T21.25 group corresponds to a shift in temperature of 1.25 °C. In this case, we observed an additional increase in proportionality (Supplementary Fig. 1E). The coefficient of variation of T20 plus T20.85 group is 4.71 % similar to that found in previous works that report proportionality of development. This analysis suggests that estimates of interindividual variability in the duration of development might be easily influenced and obscured by variability due to small changes in temperature such as those that can occur in many labs. When reducing temperature-dependent variability, the stage of postembryonic development varies independently from each other, suggesting that the duration of individual stages is regulated differentially.

### Temperature dependence of larval development

A simple method to probe developmental timing in the nematode is comparison at various temperatures. The rate of development of poikilotherms correlates with external temperature. *C. elegans* larval development is accordingly temperature dependent (Byerly et al., 1976; Filina et al., 2020). Recent studies have analyzed developmental timing at three temperatures within a limited range (15-23 °C). We therefore analyzed larval development under temperatures between 10 and 27 °C. The duration of the larval development spans from ~200 hours at 10 °C to ~36 hours at 24 °C. Above 24 °C, the duration of development lengthens (Fig. 2A). To compare the effect of temperature on each stage of development, we calculated the ratios of their duration at each temperature relative to that at 20 °C, the standard culture condition. We observed that the relative speed at extreme temperatures changed less for molts than for intermolts. The only exception is the fourth molt, which responded approximately to the same extent as intermolts (Fig. 2B).

**Figure 2.**
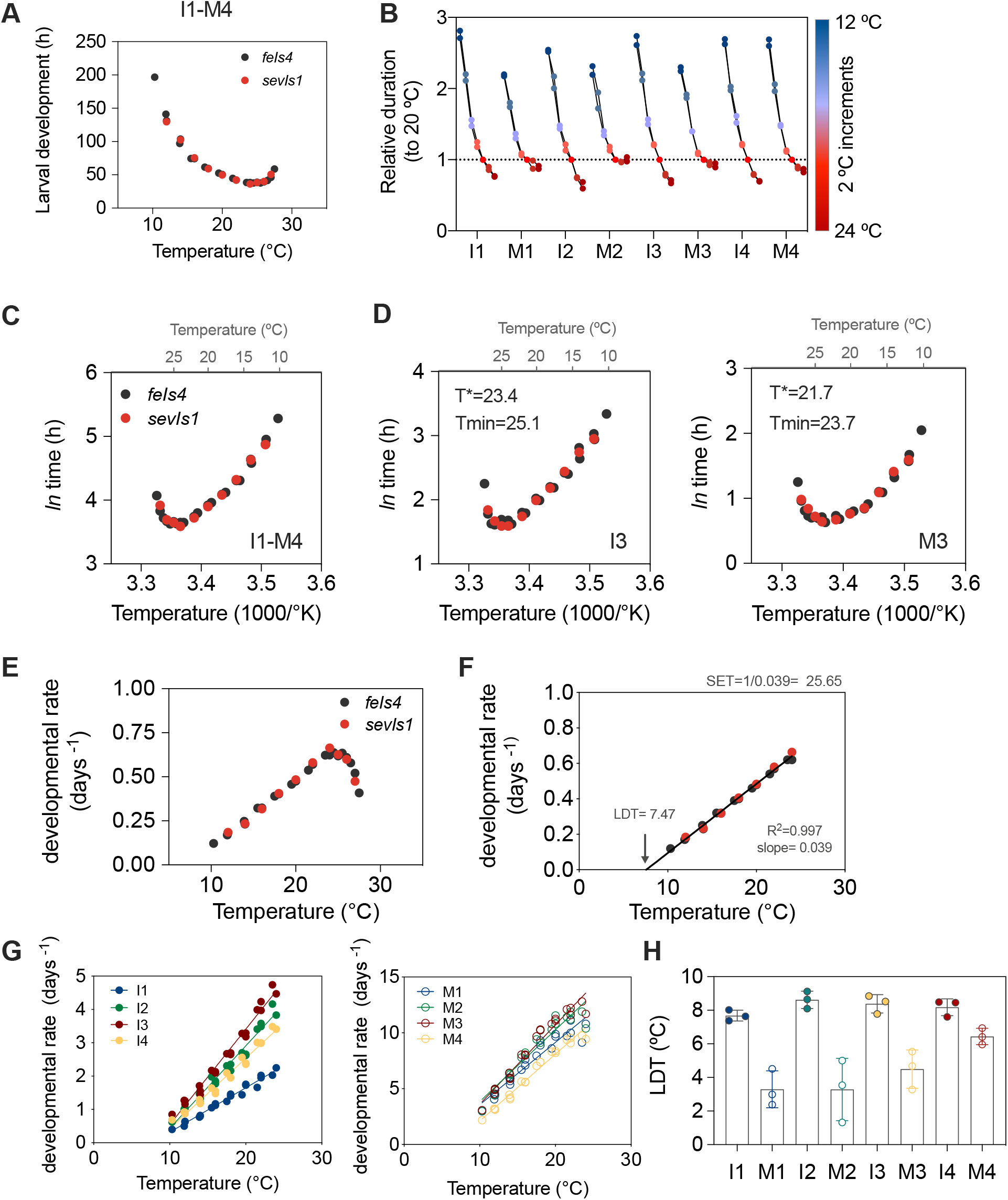
Temperature dependence of *C. elegans* postembryonic development. All panels except B represent the data for three datasets (two reporter strains/ two independent replicates; see methods section for more details). (A) Duration of larval development (I1-L4) at temperatures between 10 and 27 °C. (B) Duration of each stage relative to the duration at 12 °C for the two datasets with the same incubations temperatures (Datasets 2 and 3). (C) Arrhenius plot for the duration of development (I1-M4). (D) Same representation as in (C) for two stages of development (I3 and M3). (E) Temperature dependence of developmental rate (1/days to complete larval development). (F) Calculation of LDT and SET values from the data within the linear rate of temperature dependence (10 to 24 °C). (G) Temperature dependence of the developmental rate for each stage of development. (H) Lower developmental threshold (LDT) for each stage of development.

Unlike postembryonic development, the duration of *C. elegans* embryonic development has been tested at temperatures between 4.5 and 30 °C. In that case, temperature dependence of different embryonic intervals follows the Arrhenius equation, which describes the relationship between temperature and the rate of first-order chemical reactions (Begasse et al., 2015). In order to perform the corresponding analysis for postembryonic development, we plotted the *ln* of the duration against the inverse of the temperature in Kelvin. In this representation, the data for an interval of temperature (~12 to 24 °C) becomes linear, which indicates fitting to the Arrhenius equation (Fig. 2C, Supplementary Fig. 2A). However, when we analyzed each stage independently, differences between intermolts and molts were revealed. Namely, for the duration of the molts, the interval of temperatures that follows the Arrhenius equation was markedly reduced (Fig. 2D, Supplementary Fig. 2B, and Supplementary Table 1). We calculated two critical values, T* and Tmin, for larval development (Ls) and for the individual stages (Is and Ms; Supplementary Table 1). T* is the temperature where development deviates from the Arrhenius equation, and Tmin is the temperature that produces the fastest development (Begasse et al., 2015). For larval development, T* was 22.8 °C and Tmin was 24.5 °C, lower than the values for various stages of embryonic development. The calculation of T* showed that molts deviate from Arrhenius at lower temperatures than the corresponding intermolt (~ 1 °C). Accordingly, the temperature that sustained fastest development was lower in molts than in intermolts (Fig. 2D and Supplementary Fig. 2B). In summary, Arrhenius analysis is consistent with the analysis of relative durations. The temperature dependent acceleration of development is limited lower temperatures for molts than for intermolts.

Next, we analyzed the variation of developmental rate, defined as the inverse of the duration of development in days, in response to changes in temperature. The increase of developmental rate with temperature is linear between 10 and 24 °C (Fig. 2E). This analysis, performed for temperatures below 24 °C, allows calculation of two values that define temperature dependence of development, namely the Lower Developmental Threshold (LDT) and the Sum of Effective Temperatures (SET) (Jarośík et al., 2004) (Fig. 2F). The LDT, which is the temperature below which postembryonic development stops, corresponds to the x-intercept of the linear regression. For development as a whole, the LDT is ~7.5 °C. The SET reflects the amount of heat needed for completing development and it allows calculation of the number of days necessary to complete development at any temperature. For the entire process of development this value is 25.65 day-degrees. This means that at a temperature of 20 °C, which is 12.5 °C above the LDT, development would take 2.05 days (resulting from dividing 25.65/12.5). This is 49.24 hours, very close to the timing that we measured empirically.

We then performed the same analysis for each stage of development (Fig. 2 G) and calculated SET and LDT values. The SET values roughly reflect the duration of the larval stage, as more energy is needed to complete longer stages (Supplementary Fig. 2C). LDT calculations yielded a less intuitive result, showing differences between intermolts and molts, although these were much less pronounced for the last larval stage (Fig. 2 H). This means that intermolts effectively restrict the minimal temperature necessary for postembryonic development. Furthermore, different LDTs among stages suggests departure from Developmental Rate Isomorphy (DRI), which specifies that the proportion of total developmental time spent in a particular stage does not change with temperature (Boukal et al., 2015; Jarośík et al., 2002). Since analysis of DRI requires the use of proportional data, we calculated the duration of each stage relative to the total duration of development (Supplementary Fig. 2D). When we plotted the relative durations against the temperature, we observed that they were not constant, suggesting non-isomorphic responses between stages of development. Molts and intermolts showed a different trend (Supplementary Fig. 2E). The slope of the relative durations indicates that the fraction of development spent in molting increased at higher temperatures, while the fraction of development occupied by intermolts decreased at high temperatures. The deviations from isomorphy are larger at early stages of development, with the exception of I1 (Supplementary Fig. 2F). Ubiquity of intraspecific DRI has been questioned recently (Boukal et al., 2015; Folguera et al., 2010) and our finding of different LDT for molts and intermolts also argues against DRI in *C. elegans* postembryonic development.

Altogether, three different types of analysis suggest that, despite superficial proportionality, developmental stages differ in their response to temperature. Furthermore, since we tested a wide range of temperatures with high resolution (1-2 °C), we have been able to unveil relevant characteristics of development, such as the stage specific adherence to and deviation from the Arrhenius equation, the calculation of the temperature that sustains fastest growth, and prediction of the minimal temperature that allows larval development.

### Food quantity and quality have stage specific effects

We hypothesized that other perturbations would also affect different stages differently. Food quantity and quality affect the overall duration of *C. elegans* postembryonic development, but a detailed characterization of the effect on molts and intermolts has never been performed. We titrated the amount of the standard OP50-1 *E. coli* diet provided to the larvae and measured developmental progression. Reduction of the OP50-1 concentration from 10 g/l to 0.63 g/l did not have any effect on the duration of complete development or on the individual larval stages. However, below from the concentration of 0.31 g/l of *E. coli* OP50-1, larval development started to show a significantly increased duration. At 0.08 g/l a reduced fraction of animals (5/35) reached adulthood (Fig. 3A-B). We checked whether the duration of all the stages was reduced proportionally by calculating the ratios between the duration at each concentration relative to that at the highest concentration (10 g/l). The reduction of food to 0.16 g/l had a greater impact in the duration of I2, I3 and I4, each showing a larger than 2-fold increase in duration compared to the concentrated food source. I1, M2, M3 and M4 experienced around a 1.5-fold increase in duration in the same condition and M1 showed little variation between concentrations (Fig. 3B).

**Figure 3.**
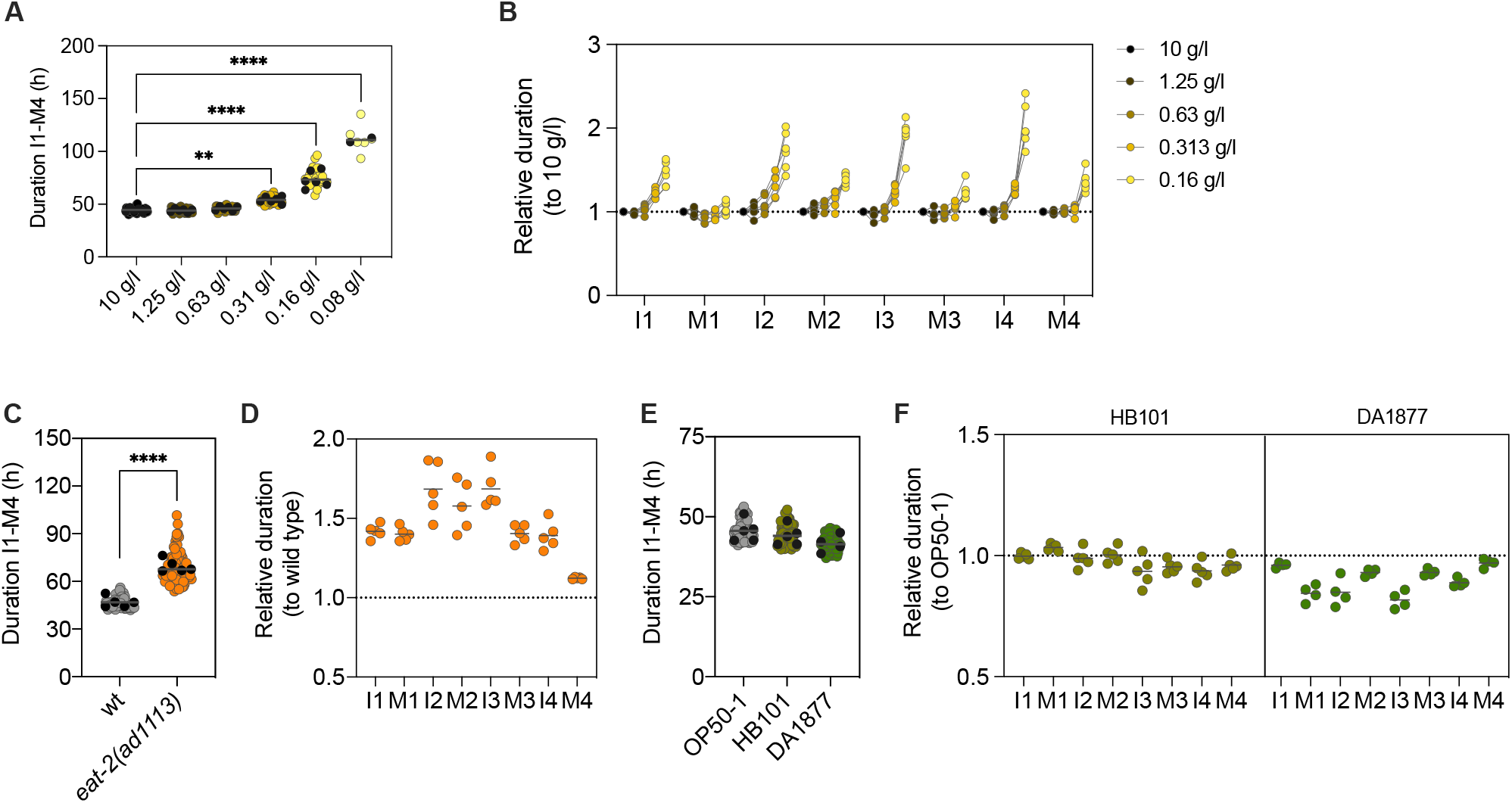
Effect of food quantity and quality in the duration of development. (A) Total duration of development at different concentrations of *E. coli* OP50-1, from 10 g/l to 0.08 g/l. (B) Duration of each stage of development relative to that at the highest concentration of food. (C) Duration of development of the wild-type strain and the *eat-2(ad1113)* mutant. (D) Duration of each stage of development of the *eat-2* mutant relative to that of the wild-type. (E) Duration of development of the wild-type strain on HB101 and DA1877 diets. (F) Duration of each stage of development on HB101 and DA1877 diets relative to the duration on OP50-1. In A, C and E, Black dots represent the mean of each experiment.

We also tested development of animals carrying the *eat-2(ad1113)* allele. *eat-2* mutants have reduced pumping rates compared to the wild-type strain and are commonly used as a model for dietary restriction (Avery, 1993; Lakowski and Hekimi, 1998). The duration of complete development for the *eat-2(ad1113)* mutant was significantly increased compared to that of the wild type (Fig. 3C). A detailed analysis of the duration of each stage in the mutant relative to wildtype showed that I2, M2, and I3 are the stages most affected by the reduction of pumping, while the duration of M4 is very similar to that of the wild type (Fig. 3D).

Food quality also alters the duration of *C. elegans* development. We used two bacterial strains that are considered good food sources for *C. elegans* (Shtonda and Avery, 2006), *E. coli* HB101 and *Comamonas* DA1877. The effect of the *Commamonas* diet is mediated by vitamin B12, the levels of which are much higher in this bacteria than in *E. coli* OP50 (Watson et al., 2014). While HB101 had only a mild, non-significant effect in the duration of complete development, the DA1877 diet reduced the duration of development by about three hours (Fig. 3E). Stage by stage analysis reveals that the duration of the stages is not reduced proportionally among stages (Fig. 3F).

Molts vary less than intermolts, both in response to temperature changes and to differences in food quality, with the exception that M4 that shows the largest temperature sensitivity among the molts (Fig. 2B and Fig. 3B). These results show that food-related interventions have different effects than do changes in temperature. Different food regimens are also distinct relative to each other. For instance, M1 is affected similarly to I1 by food titration but is accelerated by the *Comamonas* diet to a larger extent than I1 (Fig. 3B,D,F) in the *eat-2(ad113)* mutation. This demonstrates that not all modulation of developmental rate by different food-related interventions is equal. Here, vitamin B12 from the *Comamonas* diet seems to impact preferentially processes in M1, I2 and I3. In summary, each intervention shows a stage-dependent signature in their modulation of developmental rate.

### Reduced insulin signaling

The mutant *daf-2(e1370)* is commonly used as a model for reduced Insulin/IGF-1 signaling (IIS). The insulin signaling pathway promotes growth in the presence of nutrients. IIS is highly conserved and is a prominent, determinant regulator of aging and lifespan in worms, flies and mammals, including humans (Kenyon, 2010). DAF-2 is the *C. elegans* Insulin receptor, whose activation triggers a phosphorylation cascade that eventually inhibits the entrance of the transcription factor DAF-16 to the nucleus. Low signaling through the IIS pathways reduces DAF-16 phosphorylation, allowing its translocation to the nucleus, where it activates transcription of target genes. The DAF-2 receptor is activated by the binding of agonist Insulin-like peptides (ILPs) produced in response to the presence of food (reviewed in (Murphy, 2013)).

The allele *daf-2(e1370)* shows developmental delays, especially at the L2 stage (Olmedo et al., 2015; Ruaud et al., 2011). Furthermore, *daf-2(e1370)* is widely described as a thermosensitive allele. *daf-2(e1370)* forms dauer larvae at 25 °C but not at 20 °C or 15 °C. However, other phenotypes, such as lifespan extension, are present at 15 °C, suggesting that *e1370* is not thermosensitive in the classical sense but that the dauer phenotype is temperature dependent (reviewed in (Ewald et al., 2017)).

We thus decided to investigate temperature sensitivity of the developmental delay phenotype of *daf-2(e1370).* This experiment provides an outstanding opportunity to assess the combined effect of both temperature and food sensing. We tested *daf-2(e1370), daf-16(mu86)*, and the double mutant *daf-2(e1370);daf-16(mu86)* at 12, 16, 20 and 22 °C. The developmental rate of the *daf-2(e1370)* increases between 12 and 16 °C, being indistinguishable from that of the wild type. We observed no increase, however, in developmental rate between 16 and 20 °C, nor at 22.5 °C (Fig. 4A). The *daf-16* and *daf-2;daf-16* mutants developed as wild type animals at all temperatures, indicating that DAF-16 is necessary for the *daf-2* phenotype (Fig. 4A). This might suggest that *daf-2* mutants are insensitive to temperature changes below 16°C but we know that the developmental delay of *daf-2* is stage dependent at 20 °C (Olmedo et al., 2015; Ruaud et al., 2011) (Fig. 4B). We therefore analysed the effect of temperature on developmental rate of each stage independently. The developmental rate of the early stages, I1 and M1 increases in the mutant strains as in the wild type, while later larval stages increased to a lower extent in the *daf-2* mutant relative to the wild type. The most extreme difference in phenotype is found in the I2 stage, where developmental rate of the *daf-2* mutant was reduced between 16 and 20 °C. Furthermore, for L2 and L3, molts are less affected by the mutation than intermolts (Fig. 4C). Again, these phenotypes are DAF-16 dependent, as the double *daf-2;daf-16* mutants show the same developmental rate as the wild type. We also noticed temperature dependent differences in the progression of development of the mutants. At 12 °C, the successful completion of larval development required DAF-16, while at 22.5 °C, activation of DAF-16 in the *daf-2(e1370)* mutant prevented the progression of development at different stages (Supplementary Fig. 3).

**Figure 4.**
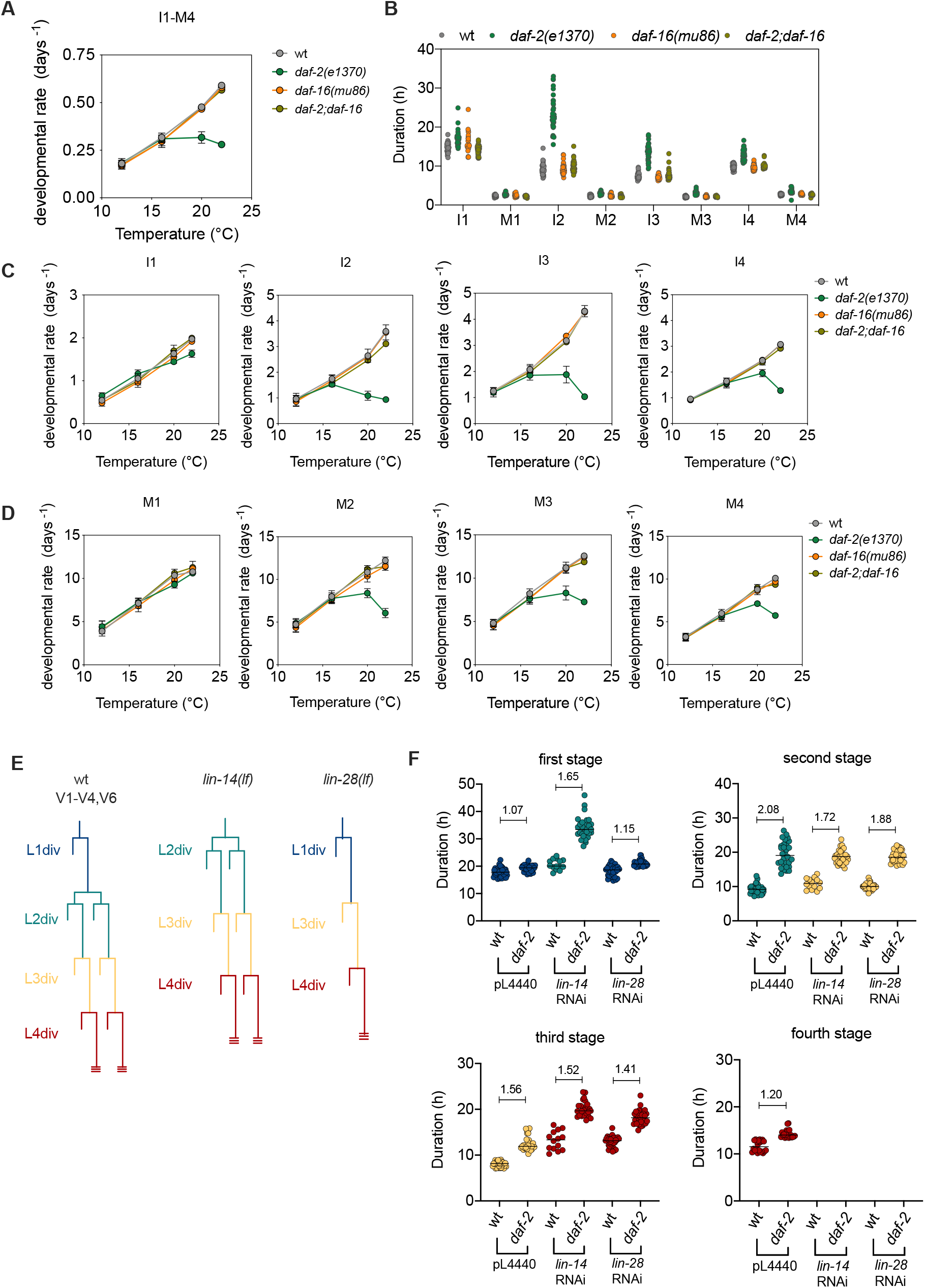
Temperature dependence of the *daf-2(e1370)* developmental rate. (A) Developmental rate of the wild type and the mutants *daf-2(e1370), daf-16(mu86)* and *daf-2(e1370);daf-16(mu86)* at 12, 16, 20 and 22 °C. (B) Duration of each stage of development at 20 °C for the four strains. Developmental rate of intermolts (C) and molts (D) at 12, 16, 20 and 22 °C, for the wild-type and mutant strains. (E) Diagram of V1-V4,V6 seam cell divisions in the wild-type strain and *lin-14* and *lin-28* lack-of-function mutants. (F) Duration of each stage of larval development. The stages are plotted as first, second, third or fourth according regardless of the cell fates expressed in the stage, which are indicated by the color of the dots, as shown in (E). The numbers inside the plots show the *daf-2* to wt ratio of the duration of each stage.

The overall developmental delay in these mutants could be explained by thermosensitivity of the *e1370* mutation, as the phenotype is only observed at high temperatures. However, the stage-dependent effect of the *daf-2(e1370)* mutation can only be explained by a different contribution of the DAF-2 receptor (via DAF-16 transcription factor function) to the progression of each stage of development.

Our results suggest that lower IIS *per se* does not lead to developmental delays. Rather, in some larval stages, low IIS and temperatures above 20 °C leads to an activation of DAF-16 that extends the duration of the stage. Stage specific extension could be the consequence of the activation of alternative developmental programs. The extended L2 observed in *daf-2(e1370)* confers resistance to the nicotinic acetylcholine receptor agonist DMPP (dimethylphenylpiperazinium) in a DAF-16 dependent manner. However, analysis of DMPP resistance of multiple *daf-2* alleles revealed that this phenotype does not correlate with their dauer phenotype (Ruaud et al., 2011). This means that the developmental program can be impacted at different stages, altering the progression of developmental events.

The duration of the L1 stage is especially insensitive to low insulin signaling. This could be a consequence of this stage being the first after hatching. Since the effect of maternal provisioning to the embryos is strongest at L1 (Supplementary Fig. 1 E), it could be that this provisioning in the newly hatched larvae is sufficient to maintain the wild-type L1 duration. Alternatively, it is also possible that the specific cellular events taking place during L1 are independent of insulin signaling. To investigate this, we knocked down the heterochronic genes *lin-14* and *lin-28* using RNAi during the development of the *daf-2(e1370)* mutant. In *lin-14* loss-of-function mutants, the blast cells that divide during postembryonic development skip the L1 specific cell fates so that hatching is followed by the L2 specific divisions. *lin-28* mutants show normal L1 divisions but premature activation of the L3 fates, skipping the L2 divisions (Fig. 4E) (Ambros and Horvitz, 1987; 1984). In the previous experiments, the *daf-2(e1370)* mutation provoked an extension of L2, L3 and L4, but not L1. When we treated *daf-2(e1370)* larvae with *lin-14* RNAi, the first observable stage, corresponding to the L2 cell fates, shows a significant extension. Treatment with *lin-28* RNAi, however, did not alter the duration of the first stage, which corresponds to the L1 divisions, as in the control (Fig. 4E). The *daf-2* mutation provoked an extension of the stages corresponding to L3 and L4 divisions in both *lin-14* and *lin-28* RNAi treatments, as occurs when these genes are knocked down in the wild type background. Therefore, our results suggest that specifically the L1 divisions are insensitive to low insulin signaling, since *lin-14* RNAi treatment, which suppresses L1 divisions, leads to strong extension of the first stage after hatching. This result supports that the duration of different larval stages can be defined by the cellular events taking place during that stage.

In addition to the marked repetitive nature of the processes, each developmental stage features specific events. While L1 is dominated by division of neuronal lineages, L2 has a key role in determination of progression to dauer or L3 larva, the L3 stage is marked by development of the somatic gonad, and L4 by initiation of meiosis of germ cells that differentiate into mature sperm. This are only some of the stage-specific events during larval development but, with this in mind, it seems remarkable that the duration of the stages is so similar. A recent work has revealed coupling between growth rate and developmental timing (Towbin and Großhans, 2021). The authors suggest two possible non-exclusive mechanisms to achieve this coupling. First, growth rate could impact a developmental clock. As a second mechanism, molting could be triggered by mechanical forces provoked by stretching of the cuticle, as also suggested recently (Nyaanga et al., 2021). This mechanism provides a plausible explanation for the similar duration of the stages that is independent of a developmental clock. However, it is difficult to infer how rate limiting interventions, as those we have applied, would impact growth rate in a stage-specific manner. Furthermore, some recent observations seem to challenge this connection between body size and developmental timing, as growth-arrested *lin-42* mutants continued development with similar dynamics to that shown before they cease growing in size (Filina et al., 2020). Rather, our results suggest that stagespecific events confer a different duration to each stage.

Altogether, our results provide strong evidence that the duration of each stage of postembryonic development is differentially affected by changes in environmental conditions, showing that their duration is differentially regulated. This regulation is compatible with a model of development based on independent linear timers for each stage of development.

## MATERIALS AND METHODS

### Culture conditions and strains

We cultured stock animals according to standard methods (Brenner, 1974), maintaining them at 20 °C on nematode growth medium (NGM) with a lawn of *Escherichia coli* OP50-1. The only exception was one of the datasets performed to analyze the effect of temperature (PE254 *feIs4*, Dataset 1), for which the animals were maintain to 18 °C prior to the analysis of development at different temperatures. A detailed description of the strains used in this study is included in Supplementary Table 2. For the experiments where we altered the food source or performed RNAi treatment by feeding, we used the bacterial strains detailed in Supplementary Table 2.

### Luminometry of single worms

We measured developmental timing using a bioluminescence-based method (Olmedo et al., 2015). For the experiments initiated with embryos, we first obtained age matched embryos by transferring 10-15 gravid hermaphrodites to a fresh NGM plate and allowing them to lay eggs over a period of approximately 2 hours. Then, using an eyelash we transferred individual embryos to the wells of a white 96-well plate containing 100 μl of S-basal (including 5 μg/ml cholesterol) with 200 μM Luciferin. For the experiment initiated from arrested L1 (Figure 2, strain PE254 *feIs4*, Dataset 1), we treated gravid adults with alkaline hypochlorite solution to obtain embryos and adjusted the concentration 20 embryos/μl of M9 buffer. The embryos were incubated overnight at 20 °C, with gentle shaking, leading to hatching and arrest at the L1 stage. Then, synchronized L1s where diluted in M9 buffer to allow pipetting of individual larvae to the wells of a white 96-well plate containing 100 μl of S-basal (including 5 μg/ml cholesterol) with 200 μM Luciferin, as before. After all embryos or larvae were placed in the wells, we added 100 μl of S-basal containing 20 g/l *E. coli* OP50-1, except when stated otherwise. We alternated the samples across the plate to avoid local effects (i.e. temperature of the luminometry reader). After preparation of the plate, we sealed the plates using a gas-permeable membrane. The plate was introduced in the luminometry reader (Berthold Centro XS3), which is placed inside a cooled incubator (Panasonic MIR-254) to allow temperature control. We measured luminesce for 1 sec at 5-min intevals, until animals reached adulthood.

### Synchronization of mothers

To test the effect of maternal age in larval development, we obtained embryos from mothers in their first, second and third day of egg laying. We prepared synchronized populations allowing 20 gravid adults to lay eggs on NGM plates for two hours. This synchronized populations where prepared approximately 136, 112, and 88 hours before the experiment to obtain mothers in their third, second, and first day of egg laying, respectively. The embryos from these mothers were used to initiate the luminometry experiments as detailed above, to measure postembryonic development.

### Temperature regulation

We measured the temperature in the plate using a data logger Thermochron iButton DS1921G (Maxim Integrated). The temperature in the plate is ~3.5 °C higher than that set at the incubator, due to the production of heat from the luminometer. All temperatures shown in the data correspond to the temperature experienced by the larvae.

For the evaluation of the effect of temperature on larval development, the animals were shifted from the maintenance temperature to the experimental temperature at the beginning of postembryonic development.

### Preparation of bacterial cultures

All strains were first grown overnight at 37 °C, shaking in LB medium with 100 μg/ml Streptomycin. Then, we diluted 1:10 in fresh LB and incubated for an additional 2.5 hours. We transferred 100 ml of the culture to two 50-ml tubes and centrifuged 10 minutes at 4,000 g at. 4 °C. We washed the bacterial pellet with 25 ml of S-basal and centrifuged again in the same conditions. After removal of the supernatant, we weighted the wet pellet and adjusted to 20 g/l using S-basal. For the experiments with different amounts of food, we performed serial dilutions to reach the desired concentrations.

### RNAi treatments

To knockdown *lin-14* and *lin-28* we performed RNAi treatments with the corresponding clones from the Vidal library. We measured development on the second generation treated with RNAi to increase the efficiency of the treatment. To grow the first generation on NGM plates, we added 200 μl of an overnight culture of RNAi or control bacteria on NGM plates with 1 mM IPTG and 100 μg/ml Ampicillin. We let the bacterial lawn dry and incubated the plates for 5 hr at 37°C and overnight at room temperature. We transferred 10 gravid adults of the strain MRS387 or MRS434 per plate and let them lay eggs for ~1 hour before removing them from the plates. We grew them for 4 days (MRS387, wild-type background) or 5 days (MRS434, *daf-2* mutant background) at 20°C. The embryos produced by these treated animals were transferred to independent wells of a 96-well plate containing 100 μl of S-basal, to measure development as described above. The only difference was that, after all embryos or larvae were placed in the wells, we added 100 μl of S-basal containing 20 g/l of the control bacteria or the corresponding RNAi clones, 2 mM IPTG and 200 μg/ml Ampicillin. The final concentration of these components 10 g/l of bacteria, 1 mM IPTG and 100 μg/ml Ampicillin. To prepare the 20 g/l bacterial suspension we proceeded as explained above, but after incubation of the diluted culture for three hours at 37 °C, we added IPTG to 1 mM and incubated for an additional two hours. Efficiency of the *lin-14* and *lin-28* treatments was confirmed by the presence of only three larval stages in the luminometry prolife. For these two conditions, larvae with four molts were excluded from the analysis.

### Summary of experimental replicates and number of animals

Figure 1 and Supplementary Fig. 1 A, B, and C contain measurements from 103 larva in six independent experiments. Supplementary Fig. 1 D represents data from three independent experiments. The number of animals for each condition is 58 (day 1), 62 (day 2), and 48 (day 3). Supplementary Fig. 1 E, to the 103 larvae analyses in Fig. 1, we added data from 17 and 18 larvae from two additional experiments.

Figure 2 includes the datasets detailed bellow. Each dataset contains one experiment at each of the indicated temperatures.

Dataset 1: Strain PE254 *feIs4*_Replicate 1. The number of larvae per condition (temperature) is 31 (10.3 °C), 26 (11.9 °C), 24 (13.9 °C), 19 (15.5 °C), 27 (17.5 °C), 33 (19.5 °C), 33 (21.5 °C), 26 (23.5 °C), 37 (24.5 °C), 18 (25.5 °C), 22 (26.5 °C), and 37 (27.5 °C).
Dataset 2: Strain PE254 *feIs4*_Replicate 2. The number of larvae per condition (temperature) is 17 (12 °C), 14 (14 °C), 21 (16°C), 19 (18 °C), 18 (20 °C), 22 (22 °C), 15 (24 °C), 19 (25 °C), 22 (26 °C) and 21 (27 °C).
Dataset 3: Strain MRS387 *sevIs1*_Replicate 2. The number of larvae per condition (temperature) is 19 (12 °C), 18 (14 °C), 17 (16 °C), 20 (18 °C), 18 (20 °C), 16 (22 °C), 21 (24 °C), 12 (25 °C), 21 (26 °C) and 18 (27 °C).

Figure 3 A shows measurement from the following number of larvae 35, 40, 40, 33, 26, and 6 for the different amounts of food, from 10 g/l to 0.08 g/l of OP50-1. Figure 3 B shows data from the same larva, this time as relative duration. In Fig. 3 B we did not include the conditions with 0.08 g/l OP50-1 due to the low number of larvae that reached adulthood. These larvae were distributed in six independent replicates. Figure 3 C and D contain results from 67 wild type and 84 *eat-2(ad1113)* larvae, in five independent experiments.

Figure 3 E and F represent the results from five experiments for HB101 and 4 experiments for DA1877, with a total of 61 larvae for OP50-1, 61 for HB101, and 59 DA1877.

Figures 4 A-D includes data from three independent experiments at each temperature. The number of larva for each condition is: at 12 °C, 41 (wt), 46 (*daf-16*), 47 (*daf-2*) and 49 (*daf-2;daf-16*), at 16 °C, 61 (wt), 39 (*daf-16*), 61 (*daf-2*) and 49 (*daf-2;daf-16*), at 20 °C, 40 (wt), 34 (*daf-16*), 40 (*daf-2*) and 52 (*daf-2;daf-16*), and at 22 °C, 53 (wt), 57 (*daf-16*), 56 (*daf-2*) and 59 (*daf-2;daf-16*), and at 22 °C.

Figure 4 F contains measurements from 33 (wt; pL4440), 14 (wt; *lin-14* RNAi), 32 (wt; *lin-28* RNAi), 30 (*daf-2;* pL4440), 30 (*daf-2; lin-14* RNAi), and 31 (*daf-2; lin-28* RNAi) larvae, in three independent experiments.

### Data analysis and statistics

We analyzed luminometry data as previously described (Olmedo et al., 2015). Shortly, we determined the timing of the molts to calculate the duration of each stage. We first calculated the moving average of the data in a time window of 12 hours. Then, we converted the raw values of luminescence to binary using as a threshold the 75% of a 12-hr moving average. To evaluate the data for onset and offset of molting we detected the transitions in the binarized data. Transitions from 1 to 0 correspond to onset of the molt and transitions from 0 to 1 correspond to offset of the molt.

To test for differences in the duration of the complete development in Figure 3A and E, we used the One-way ANOVA followed by the Dunnett’s multiple comparisons test. For the comparison in Figure 3C we used unpaired t-test. Graphs and statistics were performed on Prism 9.

### Arrhenius analysis

We used the bootstrapping regression method to obtain critical values, T* and Tmin. This statistical approach allowed us to estimate the errors associated with the interval fit regression parameters for the complete development and the individual stages. At each stage, the exponential range of the data was fit by the Arrhenius equation:

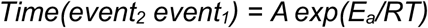

To include the high-temperature points, the equation defined by {Begasse:2015kp}, which contains an additional term to include the high-temperature data, was used:

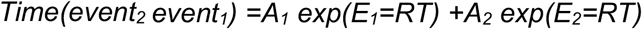

Once the terms necessary for the calculation of the Arrhenius fit were defined, the mean and SD of T* and Tmin were estimated from fits after bootstrapping with 1,000 random resamples using a custom-made script in *R* software (Supplementary File 1).

To calculate the Arrhenius interval, that is, the interval of temperatures that fit the Arrhenius equation we proceeded as in {Begasse:2015kp}. Shortly, we calculated the slope of a linear fit to the three data points centered around 20 °C. Then, we added one by one the data point from adjacent temperatures to the fit and calculated the new slopes. If the new slope shows below 10 % deviation from that of the starting interval, the added temperature is included in the Arrhenius interval.

## AUTHOR CONTRIBUTIONS

MM and MO conceived the study, AMC, FJRE, MG and MO performed experiments, AMC, FJRE, FAP, MG and MO analyzed data, all authors interpreted and discussed the results, MO wrote the manuscript, and all authors revised the manuscript.

## ACKNOWLEDGEMENTS

Some strains were provided by the *Caenorhabditis* Genetics Center (CGC), which is funded by NIH Office of Research Infrastructure Programs (P40 OD010440). We thank Jeroen van Zon (AMOLF, Amsterdam) for helpful discussion. The authors thank Sabas García-Sánchez for technical help on the experiments about the effect food on development. MO is supported by the Ramón y Cajal program of the Spanish Ministerio de Economía y Competitividad, (RYC-2014-15551). Work in the Olmedo lab is supported by the Agencia Estatal de Investigación (AEI) and the Fondo Europeo de Desarrollo Regional (FEDER) (BFU2016-74949-P, PID2019-104632GB-I00, AEI/FEDER, UE). MO and MG were supported by the Friedrich Baur Stiftung of the LMU Munich.

## SUPPLEMENTARY FIGURES AND TABLES

**Figure S1.**
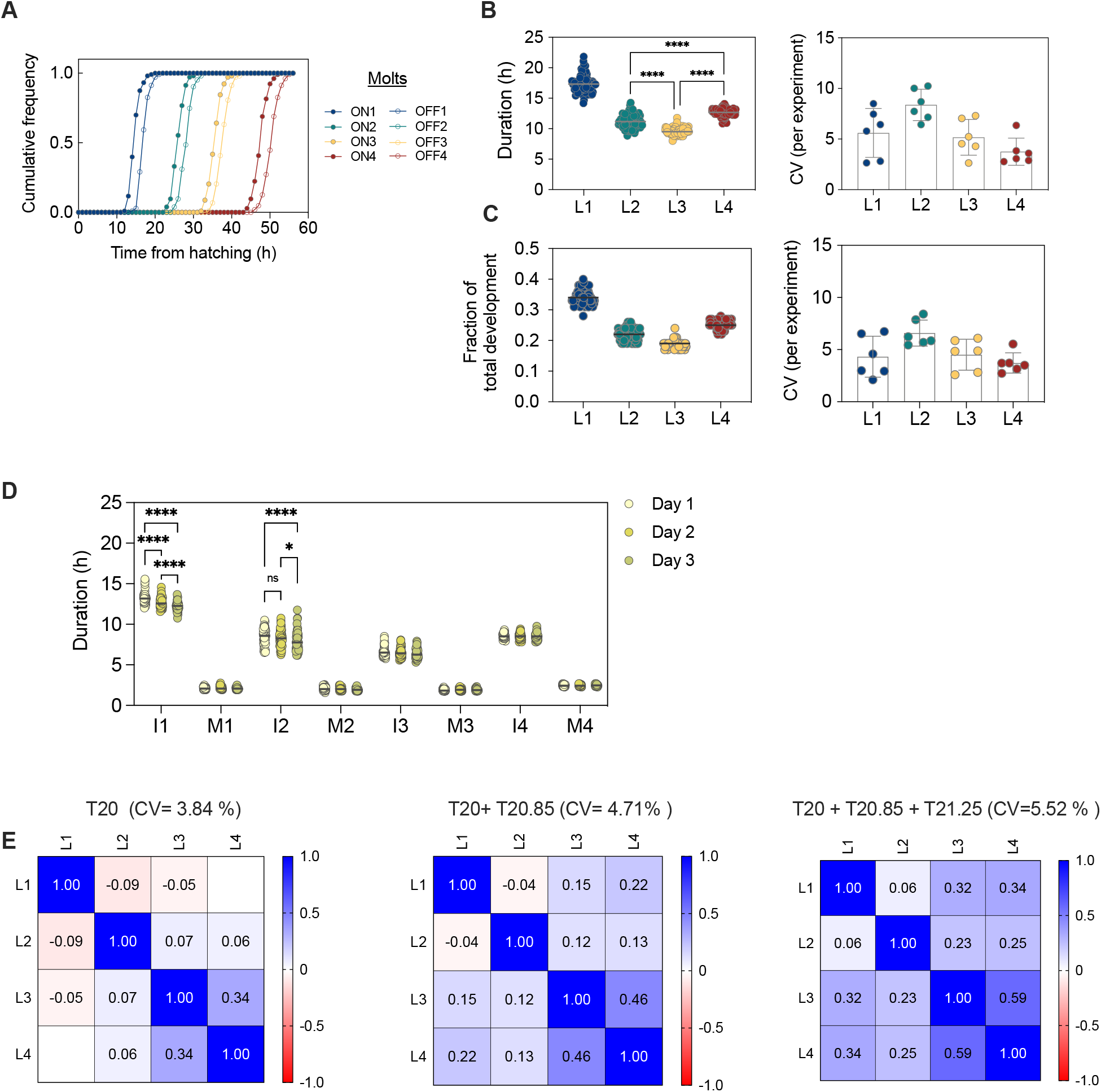
(Related to Figure 1). Interindividual variability in larval development. (A) Cumulative distribution of the beginning (ON) and end (OFF) of the four molts of postembryonic development for 103 larvae. (B) Duration and coefficient of variation (CV) for the duration of each larval stage. (C) Fraction of development dedicated to each larval stage and coefficient of variation of these values for 103 larvae. (D) Duration of each stage of development for larvae with different maternal age. Day 1, day and day 3 correspond to mothers on their first, second and third days of egg laying. (E) Correlation matrixes for each combination of larval stages for the 103 larvae in Fig. 1, and after adding 17 and 18 larvae from plates that displace the average duration by 30 min and 60 min respectively.

**Figure S2.**
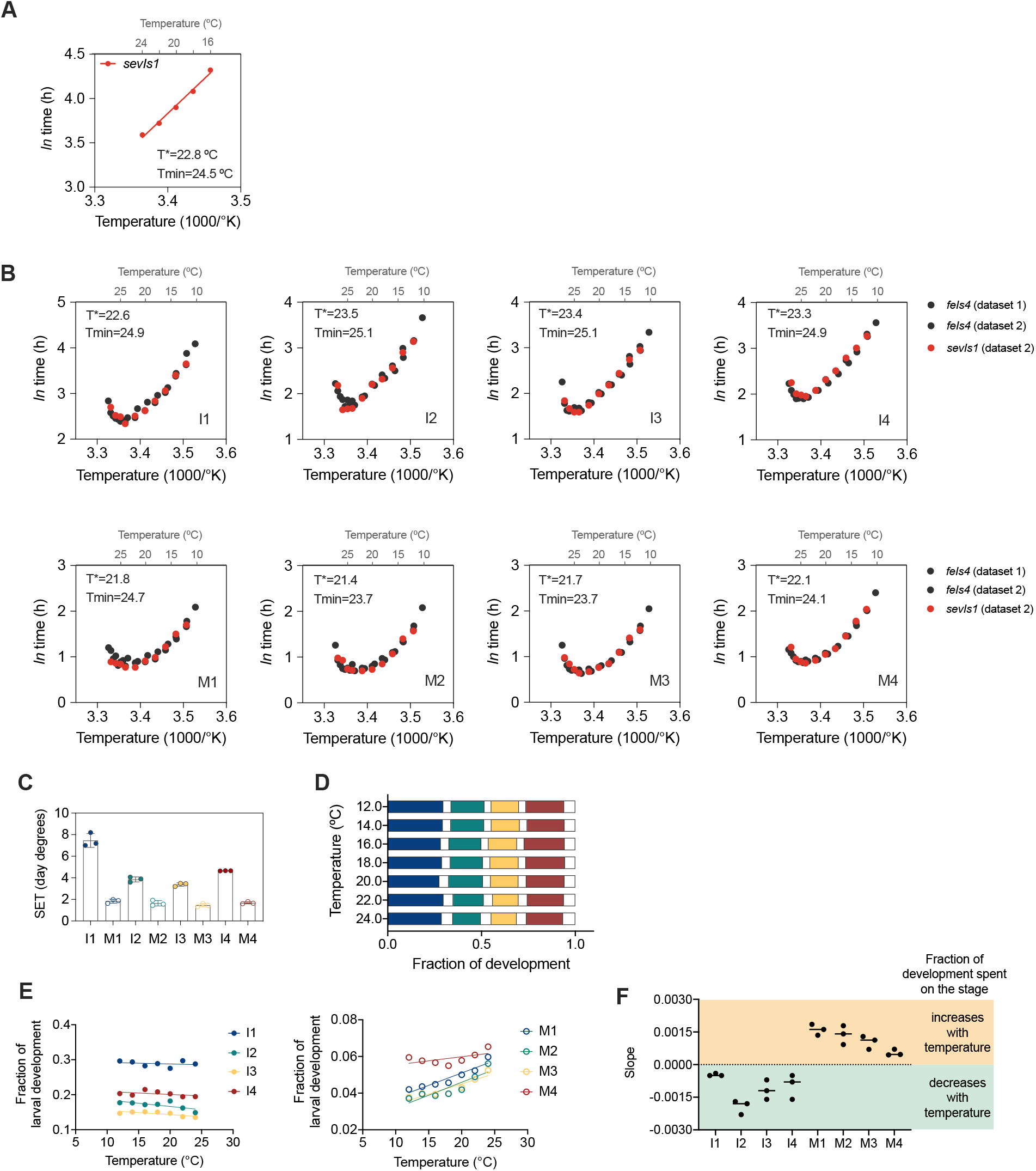
(Related to Figure 2): Temperature dependence of developmental stages. (A) Linear regression showing fitting within the Arrhenius interval of the Dataset 3. (B) Arrhenius plots for each intermolt and molt, showing the values of T* and Tmin calculated for Dataset 3. (C) SET values for each larval stage, calculated for each of the three Datasets. (D) Fraction of development devoted to each stage of development at the temperatures within the linear range defined in Fig. 2F. (E) Average fraction of development devoted to each intermolt (left) and molt (right) for Dataset 3 (F) Representative plots showing the slope of the linear regression of the fractional durations for each stage of development. The three dots for each stage correspond to each of the three datasets.

**Figure S3.**
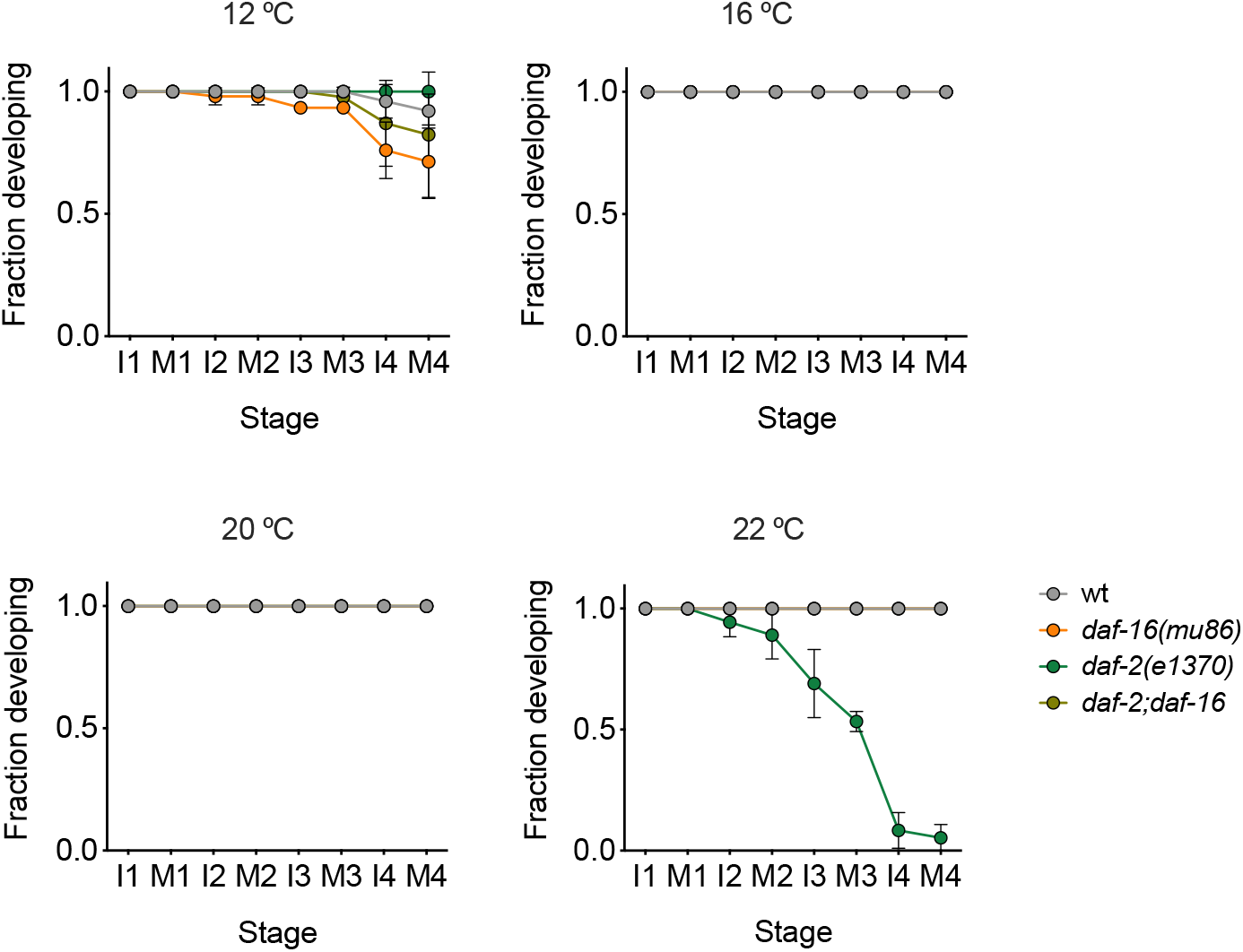
(Related to Figure 4). Fraction of animals that completes each stage of development at the different temperatures for the wild type and the *daf-2(e1370), daf-16(mu86)* and *daf-2; daf-16* mutants.

**Supplementary Table 1.** Values of T*, Tmin and Arrhenius interval for the data of the results of Dataset 3, performed with the reporter *sevIs1*. The calculation of these values is described in in the methods.

| | $T^*$ (°C) | $T_{min}$ (°C) | Arrhenius interval |
| --- | --- | --- | --- |
| **M1-M4** | $22.83 \pm 0.45$ | $24.54 \pm 0.19$ | 16 °C – 24 °C |
| **I1-M4** | $22.78 \pm 0.50$ | $24.48 \pm 0.22$ | 16 °C – 24 °C |
| **I1** | $22.63 \pm 0.93$ | $24.90 \pm 0.61$ | 16 °C – 24 °C |
| **M1** | $21.76 \pm 1.00$ | $24.69 \pm 0.64$ | 18 °C – 22 °C |
| **I2** | $23.48 \pm 0.39$ | $25.14 \pm 0.35$ | 16 °C – 26 °C |
| **M2** | $21.36 \pm 1.11$ | $23.73 \pm 0.37$ | 18 °C – 22 °C |
| **I3** | $23.37 \pm 0.68$ | $25.06 \pm 0.18$ | 16 °C – 24 °C |
| **M3** | $21.73 \pm 0.92$ | $23.74 \pm 0.33$ | 18 °C – 22 °C |
| **I4** | $23.32 \pm 0.31$ | $24.95 \pm 0.14$ | 16 °C – 22 °C |
| **M4** | $22.15 \pm 0.69$ | $24.11 \pm 0.28$ | 18 °C – 22 °C |

**Supplementary Table 2.**

| REAGENT or RESOURCE | SOURCE | IDENTIFIER |
| --- | --- | --- |
| **Bacterial Strains** |  |  |
| *E. coli*: OP50-1 | *Caenorhabditis* Genetics Center | WormBase ID: WBStrain00041971 |
| *E. coli*: HB101 | *Caenorhabditis* Genetics Center | WormBase ID: WBStrain00041075 |
| *Comamonas* sp.: DA1877 | *Caenorhabditis* Genetics Center | WormBase ID: WBStrain00040995 |
| *E. coli* HT115(DE3) pL4440 | Rual et al., 2004 | Vidal *C. elegans* RNAi feeding library |
| *E. coli* HT115(DE3) *lin-14* RNAi | Rual et al., 2004 | Vidal *C. elegans* RNAi feeding library |
| *E. coli* HT115(DE3) *lin-28* RNAi | Rual et al., 2004 | Vidal *C. elegans* RNAi feeding library |
| **Chemicals, Peptides, and Recombinant Proteins** |  |  |
| Streptomycin | Sigma-Aldrich | Cat# S6501 |
| Ampicillin | Sigma-Aldrich | Cat# A9518 |
| Tetracycline | Sigma-Aldrich | Cat# T3383 |
| IPTG | PanReac AppliChem | Cat# A4773 |
| Luciferin | BioThema | Cat# BT11-100 |
| Cholesterol | Sigma-Aldrich | Cat# C8667 |
| **Experimental Models: Organisms/Strains** |  |  |
| *C. elegans*: Strain PE254 *fels4* [*Psur-5::luc+::gfp*; *rol-6* (*su1006*)]V | Lagido et al., 2008 |  |
| *C. elegans*: Strain MRS387 *sevls1* [*Psur-5::luc+::gfp*]X | Olmedo et al., 2020 |  |
| *C. elegans*: Strain MRS424 *daf-16(mu86)I*; *sevls1* [*Psur-5::luc+::gfp*]X | Olmedo et al., 2020 |  |
| *C. elegans*: Strain MRS434 *daf-2(e1370)III*; *sevls1* [*Psur-5::luc+::gfp*]X | Olmedo et al., 2020 |  |
| *C. elegans*: MOL56 *daf-2(e1370)III*; *daf-16(mu86)I*; *sevls1* [*Psur-5::luc+::gfp*]X | Olmedo et al., 2020 |  |
| *C. elegans*: Strain MRS307 *eat-2(ad1113) II*; *fels4* [*sur-5::luc+::gfp*; *rol-6(su1006)*]V | Rodríguez-Palero et al., 2018 |  |
| **Software and Algorithms** |  |  |
| Prism 9 | GraphPad |  |
| R software | www.r-project.org |  |
| **Other** |  |  |
| Luminometry plates | Thermo | Cat# 236105 |
| Gas-permeable membrane (Breathe Easier) | SIGMA | Z763624-100EA |
| Thermochron iButton | Maxim Integrated | DS1921G |

